# *me31B* regulates stem cell homeostasis by preventing excess dedifferentiation in the *Drosophila* male germline

**DOI:** 10.1101/2021.04.07.438888

**Authors:** Lindy Jensen, Zsolt G. Venkei, George J. Watase, Bitarka Bisai, Scott Pletcher, Cheng-Yu Lee, Yukiko M. Yamashita

## Abstract

Tissue-specific stem cells maintain tissue homeostasis by providing a continuous supply of differentiated cells throughout the life of organisms. Differentiated/differentiating cells can revert back to a stem cell identity via dedifferentiation to help maintain the stem cell pool beyond the lifetime of individual stem cells. Although dedifferentiation is important to maintain the stem cell population, it is speculated to underlie tumorigenesis. Therefore, this process must be tightly controlled. Here we show that a translational regulator *me31B* plays a critical role in preventing excess dedifferentiation in the *Drosophila* male germline: in the absence of *me31B*, spermatogonia (SGs) dedifferentiate into germline stem cells (GSCs) at a dramatically elevated frequency. Our results show that the excess dedifferentiation is likely due to misregulation of *nos,* a key regulator of germ cell identity and GSC maintenance. Taken together, our data reveal negative regulation of dedifferentiation to balance stem cell maintenance with differentiation.

## Introduction

Tissue-specific adult stem cells play a critical role in sustaining tissue homeostasis by continuously providing differentiated cells throughout the life of organisms (He et al., 2009; Nystul and Spradling, 2006). The loss of stem cells or their functions underlie tissue degeneration under physiological and pathological conditions. The stem cell pool is primarily maintained by self-renewal. However, dedifferentiation, a process whereby differentiated and/or differentiating cells revert back to a stem cell identity, also helps to maintain the stem cell population beyond the lifetime of individual stem cells (de Sousa and de Sauvage, 2019; Merrell and Stanger, 2016). However, the misregulation of dedifferentiation has been implicated to underlie tumorigenesis (Landsberg et al., 2012; Schwitalla et al., 2013). Therefore, dedifferentiation must be tightly controlled to ensure stem cell maintenance, while preventing transformation. However, the molecular mechanisms that regulate dedifferentiation are not well understood.

The *Drosophila* testis serves as an excellent model system to study dedifferentiation. Notably, this model offers unambiguous identification of stem cells (germline stem cells (GSCs)) and their differentiating progeny (Fuller and Spradling, 2007; Yamashita, 2018). GSCs are attached to post-mitotic somatic hub cells, which function as a major component of the stem cell niche (Figure 1A). The hub cells secrete two major signaling ligands that promote GSC self-renewal: a cytokine-like ligand Upd that activates the JAK-STAT pathway, and a BMP ligand Dpp that activates the downstream Tkv receptor to specify stem cell identity (Kawase et al., 2004; Kiger et al., 2001; Schulz et al., 2004; Shivdasani and Ingham, 2003; Tulina and Matunis, 2001). Upon GSC divisions, daughter cells that are displaced away from the hub initiate differentiation as gonialblasts (GBs), which then continue with proliferative mitotic divisions (or transit-amplifying divisions) as spermatogonia (SGs) before entering meiotic program. SG divisions are characterized by incomplete cytokinesis, connecting all sister cells as a cluster (i.e. cyst). A membranous organelle called the fusome runs through the stabilized contractile ring, called ring canals (Figure 1A) (Yamashita, 2018).

**Figure 1.**
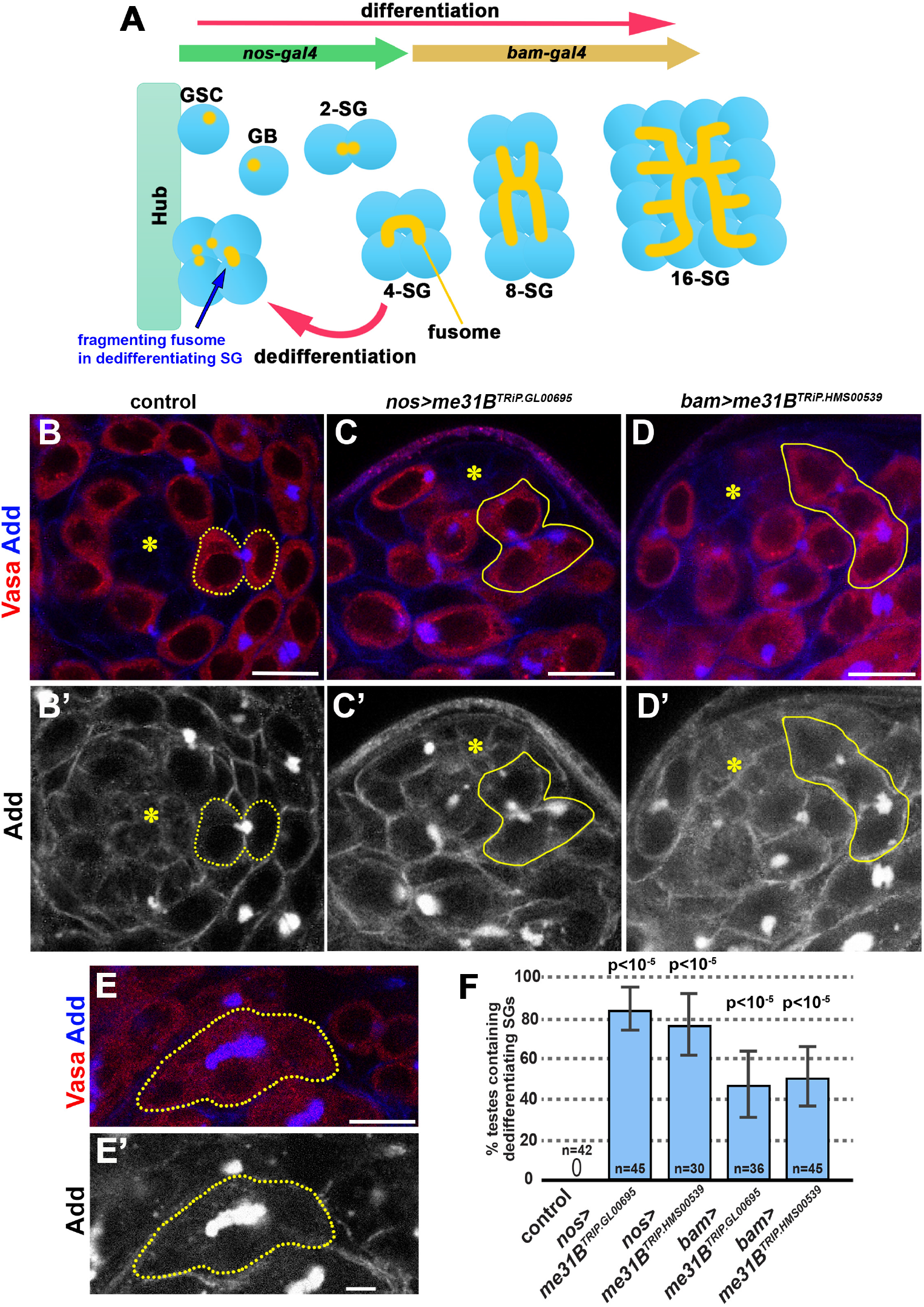
Me31B knockdown leads to excessive dedifferentiation in the *Drosophila* testis. **A**. *Drosophila* spermatogenesis. Germline stem cells (GSCs) are attached to the hub cells, which provide signaling ligands required for GSC self-renewal. Asymmetric GSC division generates a GSC and a gonialblast (GBs) that undergo 4 rounds of mitotic divisions to create 2-, 4-, 8-, and 16-cell spermatogonia (SGs). 16-cell SGs then proceed to spermatocyte stage, then to meiosis to produce sperm (not depicted). SGs can revert back to the GSC identity via dedifferentiation. During dedifferentiation, a cytoplasmic organelle called the fusome, which is normally a continuous structure that connects SGs, breaks apart. The fragmenting fusome in the dedifferentiating SG is indicated by a blue arrow. The *nos-gal4* driver is expressed in GSCs until the 4-cell SGs, whereas *bam-gal4* is expressed after the 4-cell SG stage. Note that RNAi initiated by *nos-gal4* typically perdures after *nos-gal4* expression ceases, due to persistence of RNAi (Bosch et al., 2016). **B-D.** Apical tip of the testis stained for Vasa (red, germ cells) and Adducin-like (Add, blue, fusome) in controls (B), and *nos>me31B^TRiP.GL00695^* (C), and *bam>me31B^TRiP.HMS00539^* (D) knockdown lines. Note that both RNAi lines were similarly effective, and experiments were conducted using both RNAi lines (unless the genetics crosses were too complicated to generate a desired genotype). Throughout the manuscript, examples may be shown only with one RNAi construct, but the results were confirmed by using both constructs unless otherwise noted. Yellow dotted lines indicate GSC-GB pair (B), and yellow solid lines indicate dedifferentiating SG cyst (C, D). Note that fusomes are fragmented in dedifferentiating SG cysts (C, D). Bar: 10 μm. Hub is indicated by the asterisks. **E.** An example of a continuous fusome observed in differentiating SGs (a 4-cell cyst). **F.** Frequency of testes (%) containing dedifferentiating SG cysts attached to the hub with ≥3 germ cells and fragmented fusomes in control vs. *me31B* knockdown testes. n = number of testes scored. p-value from Fisher’s exact test is provided compared to control.

Although GSCs are maintained relatively stably through consistent asymmetric divisions, which generate one GSC and one GB (Yamashita et al., 2003), GSCs can occasionally be lost (Wallenfang et al., 2006). Upon GSC loss, SGs can respond to niche vacancy, and dedifferentiate to replenish the GSC pool. During dedifferentiation of SGs, the fusome that connects SGs fragments into a more spherical structure, referred to as ‘spectrosome’ as typically observed in GSCs (Figure 1A) (Brawley and Matunis, 2004). Fragmenting fusomes in >2 cell SGs are observed only during dedifferentiation, not during differentiation, and these features can be used to unambiguously identify dedifferentiating SGs without lineage tracing (Brawley and Matunis, 2004; Sheng et al., 2009; Sheng and Matunis, 2011). Dedifferentiation was first shown in an experiment that artificially removed all GSCs via overexpression of Bam, a master regulator of differentiation (Brawley and Matunis, 2004; Sheng et al., 2009; Sheng and Matunis, 2011). While temporally controlled overexpression of Bam induced all GSCs to differentiate, withdrawal of Bam allowed SGs to repopulate the stem cell niche and produced GSCs. Subsequently, it was shown that SG dedifferentiation occurs naturally and increases during aging in unperturbed tissues (Cheng et al., 2008), suggesting that dedifferentiation is likely a mechanism that helps to maintain the GSC population throughout the lifetime of organisms, particularly with age. More recent work showed that dedifferentiation is important to sustain the GSC population under conditions that repeatedly induce GSC replenishment and challenge tissue homeostasis, such as cycles of starvation and refeeding (Herrera and Bach, 2018). SG dedifferentiation under these conditions required JNK signaling (Herrera and Bach, 2018). However, whether mechanisms exist to prevent excess dedifferentiation remain poorly understood.

*Maternally expressed at 31B (me31B)* encodes an RNA helicase of the DEAD-box family that regulates translation (Kugler et al., 2009; Kugler and Lasko, 2009; Nakamura et al., 2001). In particular, Me31B silences the translation of oocyte-localizing mRNAs, such as *oskar,* in nurse cells prior to their transport to the oocyte (McDermott et al., 2012; Nakamura et al., 2001). Me31B was also shown to repress translation of *nanos (nos*) (Gotze et al., 2017; Jeske et al., 2011), a translational regulator that is critical for germ cell specification and maintenance of GSCs (Li et al., 2009; Wang and Lin, 2004). Here, we show that *me31B* is a critical negative regulator of dedifferentiation in the *Drosophila* testis. In the absence of *me31B*, SGs frequently dedifferentiated even in the absence of known triggers, such as the induced removal of GSCs. We further show that *me31B* represses SG dedifferentiation by repressing *nos*. Our study reveals that dedifferentiation is actively repressed under normal conditions, likely to protect the native GSC population, and identifies *me31B* as a previously unknown negative regulator of dedifferentiation.

## Materials and Methods

### Fly husbandry and strains

Unless otherwise stated, all flies were raised on standard Bloomington medium at 25°, and young flies (1-to 3-day-old adults) were used for all experiments. See Supplementary Table S1 for the list of stocks used in this study.

### Immunofluorescence staining and microscopy

For Drosophila tissues, immunofluorescence staining was performed as described previously (Cheng et al., 2008). Briefly, tissues were dissected in the phosphate-buffered saline (PBS), transferred to 4% formaldehyde in PBS and fixed for 30 min. Tissues were then washed in PBS-T (PBS containing 0.1% Triton-X) for at least 30 min (three 10 min washes), followed by incubation with primary antibody in 3% bovine serum albumin (BSA) in PBS-T at 4°C overnight. Samples were washed for 60 min (three 20 min washes) in PBS-T, incubated with secondary antibody in 3% BSA in PBS-T at 4°C overnight, washed as above, and mounted in VECTASHIELD with DAPI (Vector Labs). The antibodies used are described in Supplementary Table S2. Images were taken using a Leica TCS SP8 confocal microscope with 63x oil-immersion objectives (NA = 1.4). Images were processed using Adobe Photoshop and ImageJ software. Dedifferentiating cysts were identified as a cluster of at least 3 germ cells that are clearly connected by fragmenting fusome and attached to the hub. Significance was determined using a Fischer’s Exact Test in comparison to a control.

### RNA Fluorescent in situ hybridization

To detect *nos* mRNA, single molecule fluorescent in situ hybridization (smFISH) was conducted by following a previously described protocol (Fingerhut et al., 2019). All solutions used for smFISH were RNase free. Testes from 2–3 day old flies were dissected in 1X PBS and fixed in 4% formaldehyde in 1X PBS for 30 minutes. Then testes were washed briefly in PBS before being rinsed with wash buffer (2X saline-sodium citrate (SSC), 10% formamide) and then hybridized overnight at 37°C in hybridization buffer (2X SSC, 10% dextran sulfate (sigma, D8906), 1mg/mL E. coli tRNA (sigma, R8759), 2mM Vanadyl Ribonucleoside complex (NEB S142), 0.5% BSA (Ambion, AM2618), 10% formamide). Following hybridization, samples were washed three times in wash buffer for 20 minutes each at 37°C and mounted in VECTASHIELD with DAPI (Vector Labs). Images were acquired using an upright Leica TCS SP8 confocal microscope with a 63X oil immersion objective lens (NA = 1.4) and processed using Adobe Photoshop and ImageJ software. Fluorescently labeled probes were added to the hybridization buffer to a final concentration of 50nM (for satellite DNA transcript targeted probes). Probe set against *nos* exons was designed using the Stellaris^®^ RNA FISH Probe Designer (Biosearch Technologies, Inc.) available online at https://www.biosearchtech.com/stellarisdesigner. The Stellaris^®^ RNA FISH (Biosearch Technologies, Inc.) probes were labeled with Quasar 670. Probe set was added to the hybridization buffer in 50nM final concentration. For smFISH probe sequences see Supplementary Table S3.

### RNA immunoprecipitation (RIP)-qPCR

Samples were collected from two genotypes, a control (*nos-gal4>UAS-GFP, UAS-dpp*) and an experimental (*nos-gal4>UAS-dpp, me31B-GFP*) and processed in pairs. Dpp overexpression (*UAS-dpp*) was introduced to increase SGs in the sample. ~200 testes per sample were collected into RNAse-free PBS, frozen in liquid nitrogen after removing excess liquid, and stored at −80°C until extraction. Lysis was completed by grinding the tissue in 400 μL of lysis buffer (150 mM KCl, 20 mM HEPES pH 7.4, 1mM MgCl_2_ with 1x c0mplete™ EDTA-free Protease Inhibitor Cocktail and 1U/μl RNasin^®^ Plus RNase Inhibitor from Promega added right before the use) and incubating for 30 minutes on ice with pipetting every 10 minutes. After centrifugation at 12,000xg for 5 minutes, pelleted cell debris were discarded. At this point, a 10% pre-IP input sample was removed and saved to serve as a control. For precipitation of Me31B-GFP and control GFP, GFP-conjugated magnetic beads were prepared by incubating 10 μg of mouse anti-GFP antibodies (Fisher Scientific) with 50 μL of Protein G Dynabeads™ in 200 μL of Ab Binding and Washing Buffer (provided in the kit) for 10 min at room temperature on a rotator. After antibody conjugation, beads were magnetically separated and washed once with 200 μL of Ab Binding and Washing Buffer. The antibody-conjugated beads were then incubated with the lysate for 10 minutes at room temperature (samples tubes were tumbled end-over-end during incubation). After magnetic separation of the beads, 10% of the supernatant was taken as non-bound fraction sample. The beads were washed with the Dynabeads Protein G kit Washing Buffer 3 times, and were resuspended in TRIzol (the 10% pre-IP and 10% post-IP samples were also processed with TRIzol at this time) according to the manufacturer’s instructions. cDNA was generated using SuperScript III^®^ Reverse Transcriptase (Invitrogen) followed by qPCR using *Power* SYBR Green reagent (Applied Biosystems). 10% inputs were diluted to a 1% input before RT was run. The fold enrichment was calculated by the ΔΔCt method. First, Ct values from each IP sample were normalized to their respective 1% input for each primer (ΔCt) to account for RNA sample preparation differences.

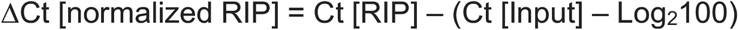

Then, the ΔΔCt (Me31B-GFP/control GFP) was obtained to compare these normalized values between the Me31B-GFP sample versus the UAS-GFP control for each primer set.

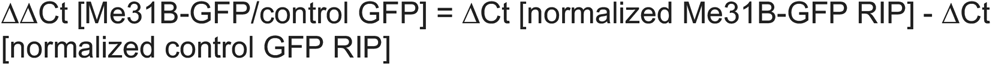

Finally, the fold enrichment was obtained by the following formula.

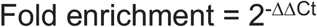

Experiments were done in technical triplicates with three biological replicates. Primers used are the following: *rp49*, forward 5’-TACAGGCCCAAGATCGTGAA-3’, reverse 5’-TCTCCTTGCGCTTCTTGGA-3’. *nanos* set #1, forward 5’-CAGTACCACTACCACTTGCTG-3’, reverse 5’-AAAGATTTTCAAGGATCGCGC-3’. Nanos set #2, forward 5’-CACCGCCAATTCGCTCCTTAT-3’, reverse 5’-GCTGGTGACTCGCACTAGC-3’. *bam*, forward 5’-TGACGTTACTGCACCACTCC-3’, reverse 5’-CGAACAGATAGTCCGAGGGC-3’.

## Results

### *me31B* prevents excess dedifferentiation of SGs in *Drosophila* testes

To study the role of *me31B* in the testis, we used two independent RNAi constructs (*UAS-me31B^TRiP.GL00695^* and *UAS-me31B^TRiP.HMS00539^*, available from Bloomington Stock Center, see methods). Using these constructs and the *nos-gal4* driver, we knocked down *me31B* in germ cells (Supplementary Figure S1, *nos-gal4>UAS-me31B^TRiP.GL00695^* and *nos-gal4>UAS-me31B^TRiP.HMS00539^*, hereafter *nos>me31B^TRiP.GL00695^* and *nos>me31B^TRiP.HMS00539^*, respectively, or simply *nos>me31B^RNAi^* as essentially the same results were obtained with both RNAi constructs). We found that Me31B-GFP was expressed in both germline and somatic cells in the testis, and the GFP signal was substantially reduced in the germline upon expression of the RNAi construct using *nos-gal4*, confirming the efficiency of these RNAi constructs (Supplementary Figure S1). Although Me31B has been reported to be a component of nuage (germ granules) (DeHaan et al., 2017; Liu et al., 2011; Thomson et al., 2008), we observed diffuse cytoplasmic localization of Me31B-GFP in germ cells in the adult testis and Me31B-GFP did not co-localize with the nuage marker Vasa in control flies. Moreover, *me31B* knockdown did not affect nuage morphology (Supplementary Figure S1).

As expected, GSCs in control testes surrounded the hub and were either single cells or connected to their immediate daughter cells (GBs) prior to completion of cytokinesis (Figure 1B). Intriguingly, *nos>me31B^RNAi^* testes often contained SG cysts that were attached to the hub cells, as opposed to control testes where only single cells (GSCs) or doublets (GSC-GB pairs) were attached to the hub (Figure 1B). Their identity as SG cysts is based on the fact that they contained ≥3 germ cells that were connected to each other (Figure 1C-D). The fusomes in these SG cysts at the hub in *nos>me31B^RNAi^* testes were fragmented (Figure 1C-D), a well-established hallmark of dedifferentiating SGs (Brawley and Matunis, 2004; Sheng et al., 2009; Sheng and Matunis, 2011), rather than continuous as in differentiating SGs (Figure 1E). We observed dedifferentiating SG cysts, identified by their fragmented fusomes and attachment to the hub, in about 80% of *nos>me31B^RNAi^* testes but not in any control testes (Figure 1F). The number of SGs within dedifferentiating SG cysts was not always 2^n^: often they contained 3 SGs, indicating that some SGs might have already dedifferentiated into single GSCs or died during dedifferentiation.

We considered two possibilities that could explain this phenotype. First, *me31B* may be required in SGs to directly prevent their dedifferentiation. In addition, *me31B* may be required to maintain GSCs in the niche, which would indirectly prevent SG dedifferentiation. To determine if *me31B* acts directly in SGs, we used the *bam-gal4* driver to deplete *me31B* only in the 4-cell SG and later stages (Chen and McKearin, 2003b). We found that about 50% of *bam>me31B^RNAi^* testes displayed SG cysts with ≥3 germ cells attached to the hub cells and fragmented fusomes (Figure 1D, F). These results demonstrate that *me31B* is required in SGs in a cell autonomous manner to prevent their dedifferentiation; however, we note that the frequency of dedifferentiation is higher when RNAi constructs were driven by *nos-gal4* than by *bam-gal4*, suggesting that *me31B* may have additional functions in early germ cells to prevent dedifferentiation (see below).

### Dedifferentiating SGs activate BMP signaling

GSC identity in the *Drosophila* testis is specified by JAK-STAT and BMP signaling (Kawase et al., 2004; Kiger et al., 2001; Schulz et al., 2004; Shivdasani and Ingham, 2003; Tulina and Matunis, 2001). We examined whether the activation of these pathways was altered upon knockdown of *me31B*.

We found that GSCs in *bam>me31B^RNAi^* testes had similar STAT expression as control testes (Supplementary Figure S2C-D), suggesting that dedifferentiation induced in *bam>me31B^RNAi^* testes is not due to altered STAT signaling. Importantly, when a cyst of dedifferentiating *bam>me31B^RNAi^* SGs was attached to the hub cells, only the germ cells that were in direct contact with the hub had high STAT levels (Supplementary Figure S2D, arrow). Thus, our data suggest that deregulation STAT signaling cannot explain the enhanced dedifferentiation upon knockdown of *me31B*, and that *me31B* acts independently of STAT activation in SGs to prevent their dedifferentiation. Additionally, these results indicate that germ cells in ≥4-cell SG cysts can reestablish STAT signaling upon homing into the niche during dedifferentiation triggered by depletion of *me31B*. Although downregulation of JAK-STAT signaling is reported to prevent SG dedifferentiation (Sheng et al., 2009), our data suggest that the dedifferentiation induced by depletion of *me31B* does not directly involve activation the JAK-STAT pathway. We speculate that JAK-STAT signaling might help maintain GSCs that were generated by dedifferentiation, instead of inducing dedifferentiation *per se*. However, STAT expression was reduced in GSCs of the *nos>me31B^RNAi^* testes compared to controls (Supplementary Figure S2A-B), suggesting that *me31B* has an additional role in GSCs to maintain STAT activation. Reduced STAT in *nos>me31B^RNAi^* testes may explain why we observe a higher frequency of dedifferentiation with *nos-gal4-*driven knockdown of *me31B* compared to *bam-gal4*-driven knockdown of *me31B* (Figure 1F).

In wild-type testes, activation of BMP signaling triggers phosphorylation of Mad (pMad) in GSCs and in GBs that are still connected to GSCs (Kawase et al., 2004) (Figure 2A). We found that knockdown of *me31B*, either by *nos-gal4* or *bam-gal4*, resulted in a high pMad signal in germ cells outside GSCs and GBs (Figure 2B, C). Moreover, in *me31B* knockdown testes, we observed high pMad signal in all the germ cells within a dedifferentiating SG cyst attached to the hub (Figure 2B, C) and even in SGs that were not yet attached to the hub (Figure 2B). We observed pMad-positive germ cells outside the niche in only 7.7% of control testis (n=39 testes), but in over 50% of *me31B* knockdown testes (91.7% in *nos*>*me31B^TRiP.HMS00539^*, n=48, 66.7% in *nos*>*me31B^TRiP.GL00695^*, n=18, 58.8% in *bam*>*me31B^TRiP.HMS00539^*, n=34, 54.8% in *bam*>*me31B^TRiP.GL00695^*, n=31). These results indicate that the activation of BMP signaling precedes the re-acquisition of GSC identity during dedifferentiation due to *me31B* depletion, and may mediate dedifferentiation. Indeed, we found that overexpression of constitutively active Tkv (Tkv*) (Nellen et al., 1996), the receptor of BMP ligands, either by *nos-gal4* or *bam-gal4,* was sufficient to induce dedifferentiation (Figure 2D). Taken together, we propose that *me31B* may prevent dedifferentiation of SGs by directly or indirectly downregulating BMP signaling.

**Figure 2.**
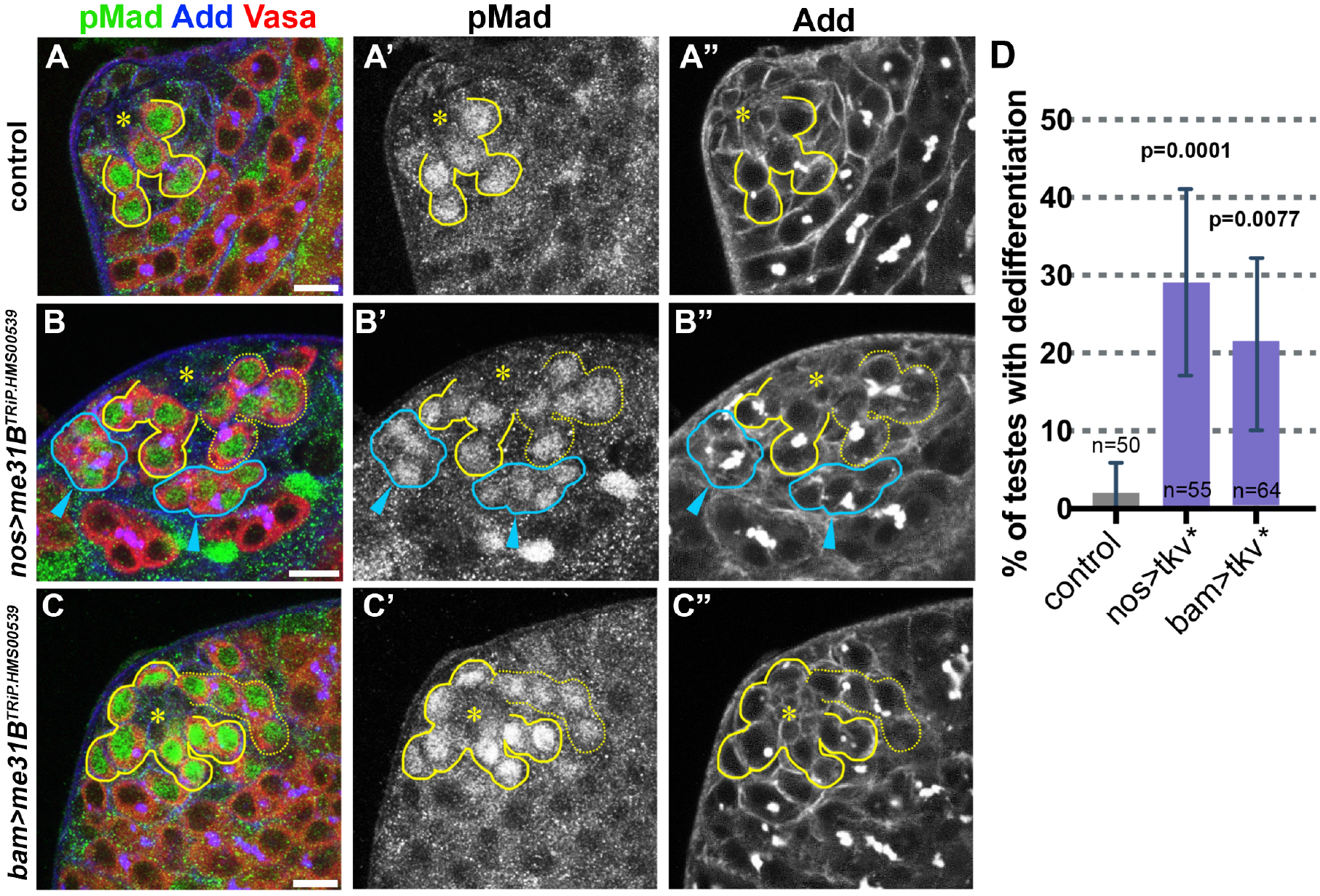
BMP signaling is upregulated upon knockdown of *me31B*. A-C. Apical tip of the testes in control (A), *nos-gal4>me31B^TRiP.HMS00539^* (B), or *bam-gal4> me31B^TRiP.HMS00539^* (C) stained for pMad (green), Vasa (red), and Adducin-like (blue). Bar: 10μm. Hub is indicated by the asterisks. GSCs and connected GBs are indicated by yellow lines. Dedifferentiating cysts that are attached to the hub are indicated by yellow dotted lines. Dedifferentiating cysts that are not yet attached to the hub are indicated by blue lines and arrowheads. D. Ectopic expression of constitutive active Tkv (Tkv*) either by *nos-gal4* driver or *bam-gal4* driver results in elevated dedifferentiation. n=number of testes scored. p-value from Fisher’s exact test is provided compared to control.

### Knockdown of *me31B* leads to misregulation of *nos* expression

Previous work showed that Me31B silences *nos* mRNA translation during embryonic development of *Drosophila* (Gotze et al., 2017; Jeske et al., 2011). In the adult germline, Nos instructs germ cell identity and GSC maintenance via translational repression of critical targets, such as Bam (Li et al., 2009; Wang and Lin, 2004) and a regulatory feedback exists between *nos*, Mad and *bam* controls germ cell differentiation (Harris et al., 2011).

To investigate whether Me31B might regulate *nos* mRNA translation during spermatogenesis, we examined Nos protein levels upon knockdown of *me31B*. In control testis, we detected Nos protein in early-stage germ cells (GSC to 4-cell stage SGs) (Figure 3A). In contrast, upon knockdown of *me31B* either by *nos-gal4* or *bam-gal4*, we observed Nos protein even in 8-cell SGs (Figure 3B, C, D), consistent with Me31B downregulating *nos* mRNA translation in the *Drosophila* testis. Nos and Bam, a master regulator of differentiation (McKearin and Ohlstein, 1995; McKearin and Spradling, 1990), are expressed in a reciprocal manner and act antagonistically in stem cell maintenance and differentiation in the *Drosophila* germline (Chen and McKearin, 2005; Li et al., 2009). Indeed, *me31B* knockdown in the testes led to delayed Bam expression and a dramatic increase in the frequency of 4-cell SGs that lacked Bam protein (Supplementary Figure S3).

**Figure 3.**
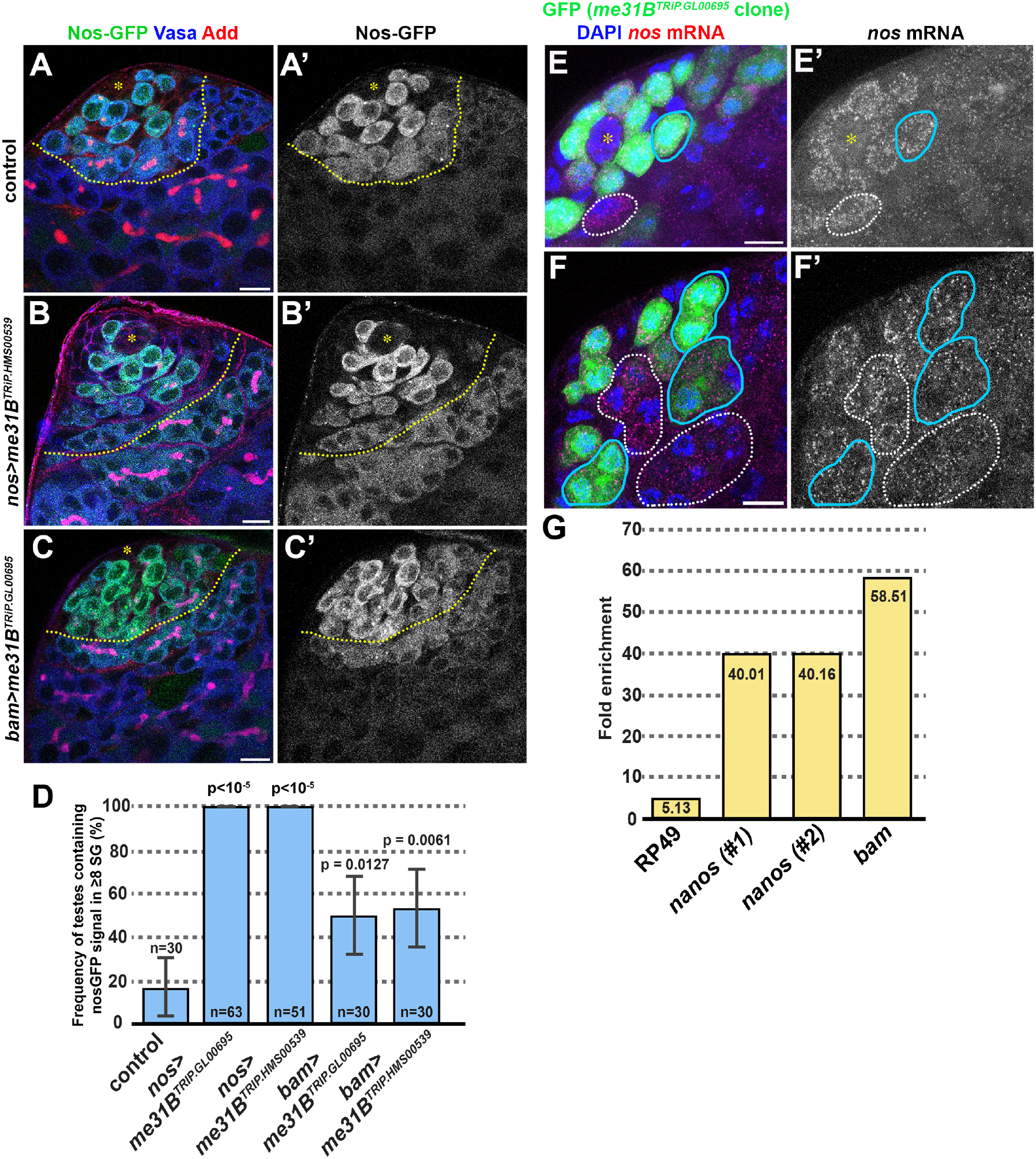
Me31B binds to *nos* and *bam* mRNA to promote SG differentiation. A-C. Apical tip of the testes expressing *nos-GFP* under the control of endogenous promoter and 3’UTR, stained for Vasa (blue) and Adducin-like (red). Control (A), *nos >me31B^TRiP.HMS00539^* (B), or *bam>me31B^TRiP.GL00695^* (C). Bar: 10μm. Hub is indicated by the asterisks. The boundary between 4-cell and 8-cell SGs is indicated by yellow dotted lines. D. The frequency of the testes that contains Nos-GFP-positive ≥8-cell SGs. n=number of testes scored. p-value from Fisher’s exact test is provided compared to control. E, F. Apical tip of the testes probed for *nos* mRNA with single molecule RNA *in situ* hybridization. GFP clones co-express *me31B^TRiP.GL00695^*, both driven by *nos-gal4*. Examples of *me31B^TRiP.GL00695^* clone cysts are indicated by blue lines, and the wild type (control) cysts are indicated by white dotted lines. (*hs-FLP, nos-FRT-stop-FRT-gal4, UAS-GFP, UAS*-*me31B^TRiP.GL00695^* flies were subjected to heat shock to activate *nos-gal4* to induce *me31B^TRiP.GL00695^* clones. See methods for details). G. Me31B-GFP RIP-qPCR probed for two sets of primers for *nos* mRNA and a primer set for *bam* mRNA, demonstrating that both *nos* mRNA and *bam* mRNA are highly enriched upon pulldown of Me31B-GFP protein.

To determine if Me31b regulates *nos* mRNA levels, we utilized a system that allows a direct comparison of germ cells with and without knockdown of *me31B* within the same tissue. Briefly, we generated testes with germ cell clones that co-express *me31B^RNAi^*and GFP (*hs-FLP, nos-FRT-stop-FRT-gal4*, *UAS-GFP*, *UAS-me31B^RNAi^*), and detected *nos* mRNA by single molecule RNA *in situ* hybridization (see methods). We compared GFP+ (*me31B^RNAi^*) vs. GFP-(control) germ cells within the same tissue and did not observe a detectable difference in the level of *nos* mRNA either in early germ cells (Figure 3E) or late SGs (Figure 3F), suggesting that *me31B* does not regulate *nos* mRNA levels. Taken together, these results suggest that Me31B regulates *nos* mRNA translation but not mRNA levels, consistent with other contexts where Me31B acts as a regulator of translation (Nakamura et al., 2001; Peter et al., 2019; Wang et al., 2017).

To determine if Me31B might regulate *nos* mRNA translation via direct binding, we performed RNA immunoprecipitation (RIP)-qPCR with testes expressing Me31B-GFP or GFP as a control (see methods). We found that *nos* mRNA co-immunoprecipitated with Me31B-GFP (Figure 3G). Interestingly, *bam* mRNA also co-immunoprecipitated with Me31B-GFP (Figure 3G), implying that Me31B may directly regulate both *nos* and *bam*. These results indicate that *nos* mRNA is a direct target of Me31B in the testis, and identify *bam* mRNA as an additional target. Overall, we conclude that *me31B* prevents dedifferentiation of SGs by reducing Nos protein levels and increasing Bam protein levels.

### *nos* is necessary and sufficient for dedifferentiation

Based on the results above, we hypothesized that Me31B prevents dedifferentiation in late SGs by silencing *nos* mRNA translation. This hypothesis predicts that *nos* downregulation would rescue the elevated dedifferentiation caused by knockdown of *me31B*. Indeed, we found that simultaneous knockdown of *nos* and *me31B* greatly reduced dedifferentiation to the level of the wild type control (Figure 4A). These data suggest that *nos* is the main functional target of *me31B* in repressing dedifferentiation. To verify that the reduced dedifferentiation in the double knockdown lines is not due to the presence of two UAS-driven transgenes and dilution of the gal4 driver, we tested a control genotype expressing *me31B^RNAi^* and a GFP transgene under the control of UAS. This genotype maintained the high frequency of dedifferentiation despite the presence of two UAS-driven transgenes (Figure 4A). Therefore, we infer that *nos* is necessary for the dedifferentiation induced by knockdown of *me31B*.

**Figure 4.**
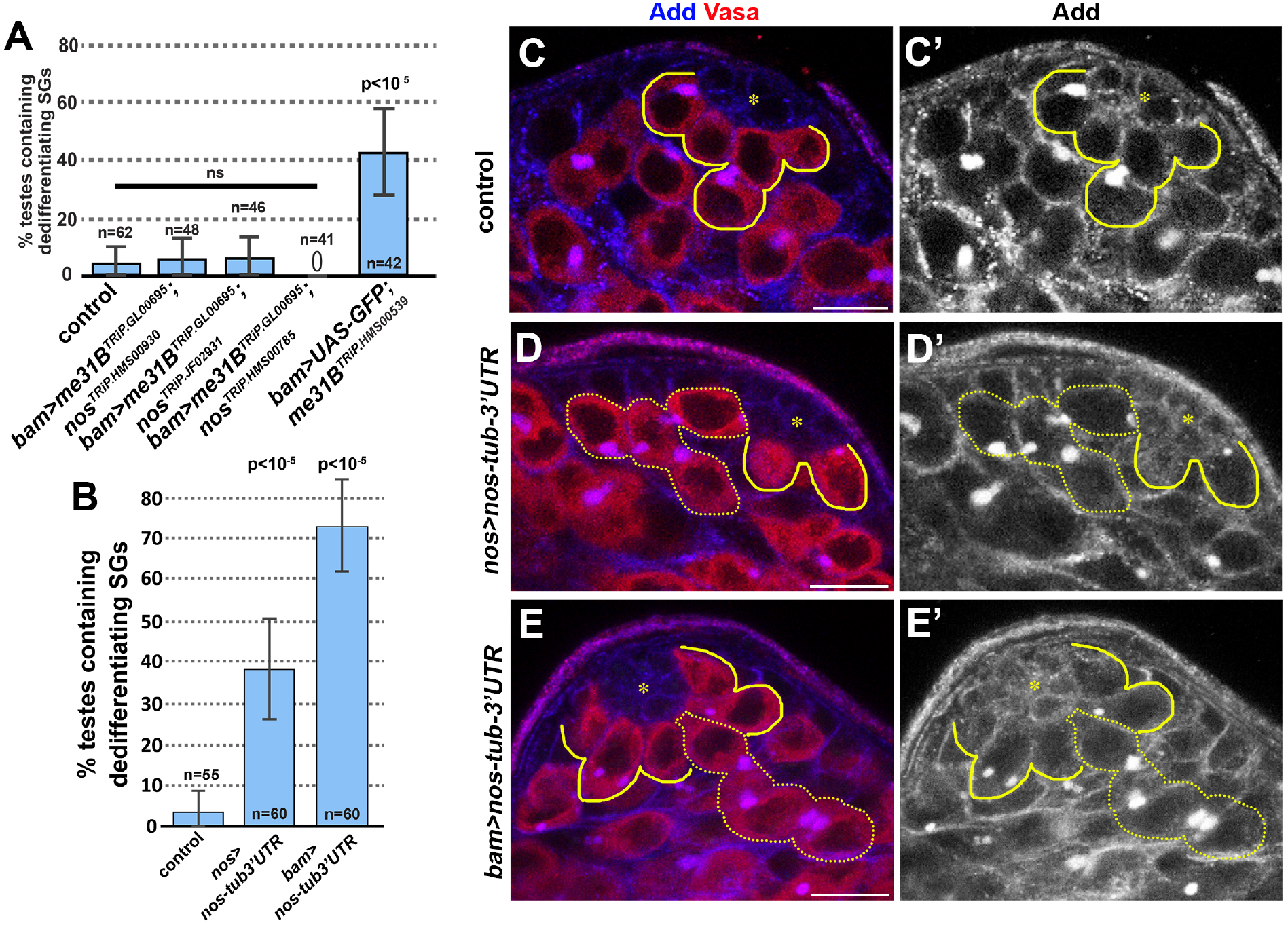
*nos* is necessary and sufficient for dedifferentiation. A. Frequency of testes containing dedifferentiating cysts in the indicated genotypes. Knockdown of *nos* diminishes dedifferentiation due to *me31B* knockdown. n=number of testes scored. p-value from Fisher’s exact test is provided compared to control. ns: not statistically significant (p>0.5) B. Frequency of testes containing dedifferentiating cysts upon ectopic expression of *nos* with *tubulin 3’UTR* (*nos-tub3’UTR*) driven by *nos-gal4* or *bam-gal4*. p-value from Fisher’s exact test is provided compared to control. C-E. Apical tip of testes from control testis (C), testis expressing *nos-tub3’UTR* by *nos-gal4* (D) or *bam-gal4* (D). GSCs and connected GBs are indicated by solid yellow lines, and dedifferentiating cysts are indicated by dotted yellow lines. Bar: 10μm. Hub is indicated by the asterisks.

Moreover, we found that upregulation of *nos* was sufficient to induce dedifferentiation. Briefly, we employed a *nos* transgene in which the 3’UTR was replaced by the *tubulin* 3’UTR (*UAS-nos-tub3’UTR*), which disrupts the regulation of *nos* by translational repressors such as Me31B (Gavis and Lehmann, 1994). When the *UAS-nos-tub3’UTR* transgene was expressed with the *nos-gal4* driver, we found that ~40% of testes contained dedifferentiating SGs, as opposed to ~3% in control (Figure 4B, C, D). Moreover, when the *UAS-nos-tub3’UTR* transgene was driven by *bam-gal4*, we observed an even higher frequency of dedifferentiation (~70%) (Figure 4B, E). These results suggest that upregulation of *nos* is sufficient to induce dedifferentiation.

Interestingly, when *me31B* knockdown was combined with *nos-tub3’UTR* expression under the control of the *nos-gal4* driver, it led to a near complete block of differentiation (*nos>nos-tub3’UTR, me31B^TRiP.HMS00539^*)(Figure 5). The differentiation block was so severe that our criteria of dedifferentiation used above (i.e. connected cells at the hub with fragmented fusomes) was not applicable: 29% of testes (n=45 testes) contained SGs but never progressed to spermatocyte differentiation (which can be recognized by growth in cell size) (Figure 5B). In addition, 91% of testes (n=45 testes) contained SG/SC cysts that contains ≥32 cells, further suggesting the failure in differentiation into spermatocyte stage (Figure 5C). These results may imply that additional targets of *me31B* cooperate with misregulated *nos* to enhance the phenotype. Alternatively, further upregulation of endogenous *nos* due to *me31B* depletion and the *nos-tub3’UTR* transgene may enhance the effect.

**Figure 5.**
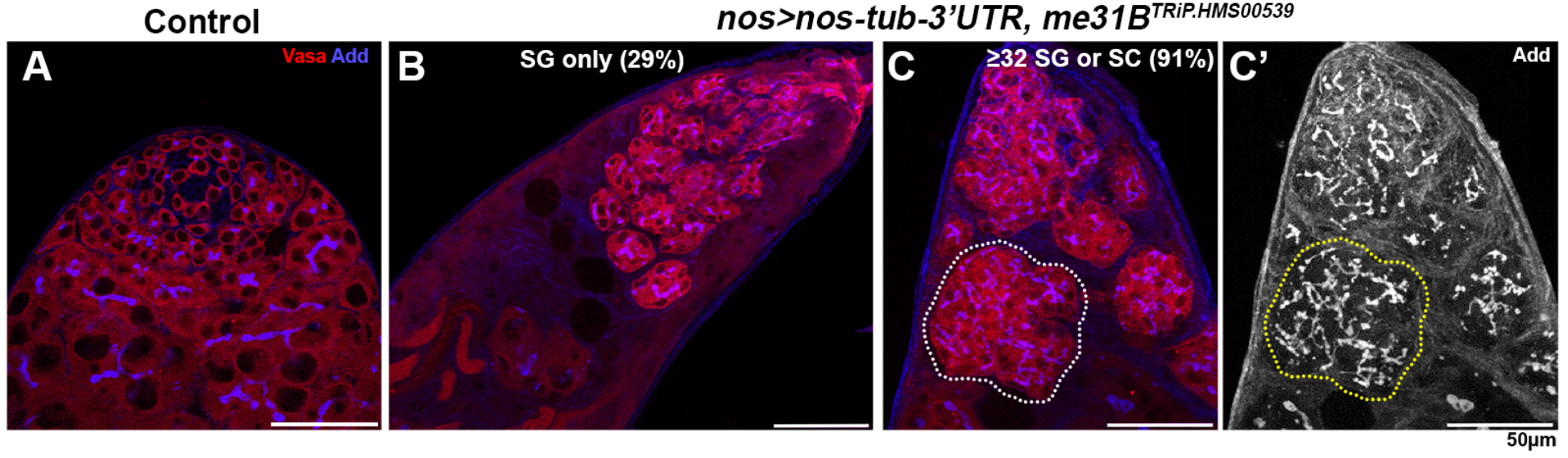
Combination of *nos* upregulation and *me31B* knockdown blocks differentiation. A. Apical tip of the testes stained for Vasa (red) and Adducin-like (blue) in control (A), or *nos>nos-tub3’UTR, me31B^TRiP.HMS00539^*(B, C). A cyst that contains ≫16 SGs is indicated by dotted lines in C. Bar: 50μm.

### Nos expression is dynamically regulated at multiple levels during differentiation in the male germline

Regulation of *nos* mRNA translation has been well documented and intensively studied, particularly in the context of germ cell specification (Gavis and Lehmann, 1992, 1994; Kugler and Lasko, 2009). The regulation of mRNA translation is critically important during oocyte development: the mRNAs that specify germ cell fate in the embryos, including *nos* and *osk* mRNA, are transcribed in nurse cells, transported into developing oocytes, and stored in mature oocytes to be translated later (Lehmann, 2016). Accordingly, mRNA synthesis (transcription) is spatially and temporally separated from protein production (translation), making it critically important to control the timing of translation by both translational repression and activation.

Whether *nos* transcription is spatiotemporally distinct from Nos protein production during the development of male germ cells in the testis is not known. To address this question, we generated a *nos* promoter reporter by driving a destabilized GFP (d2EGFP) fused to the *hsp70 3’UTR* from the *nos* promoter (Figure 6A). Because neither the mRNA nor protein products are stable in this reporter, the GFP signal closely recapitulates the activity of the promoter. Interestingly, we found that the *nos* promoter is active only in GSCs (and GBs that are still connected to GSCs (Figure 6B), suggesting that *nos* is transcribed only in these early germ cells. These data suggest that Nos protein, which is observed within GSCs through to 4-cell stage SGs, is primarily produced by translation of *nos* mRNA inherited by 2-cell SG and 4-cell SGs (Figure 6C). In addition, stable Nos protein generated in GSCs and GBs may contribute to its persistence through to the 4-cell SG stage.

**Figure 6.**
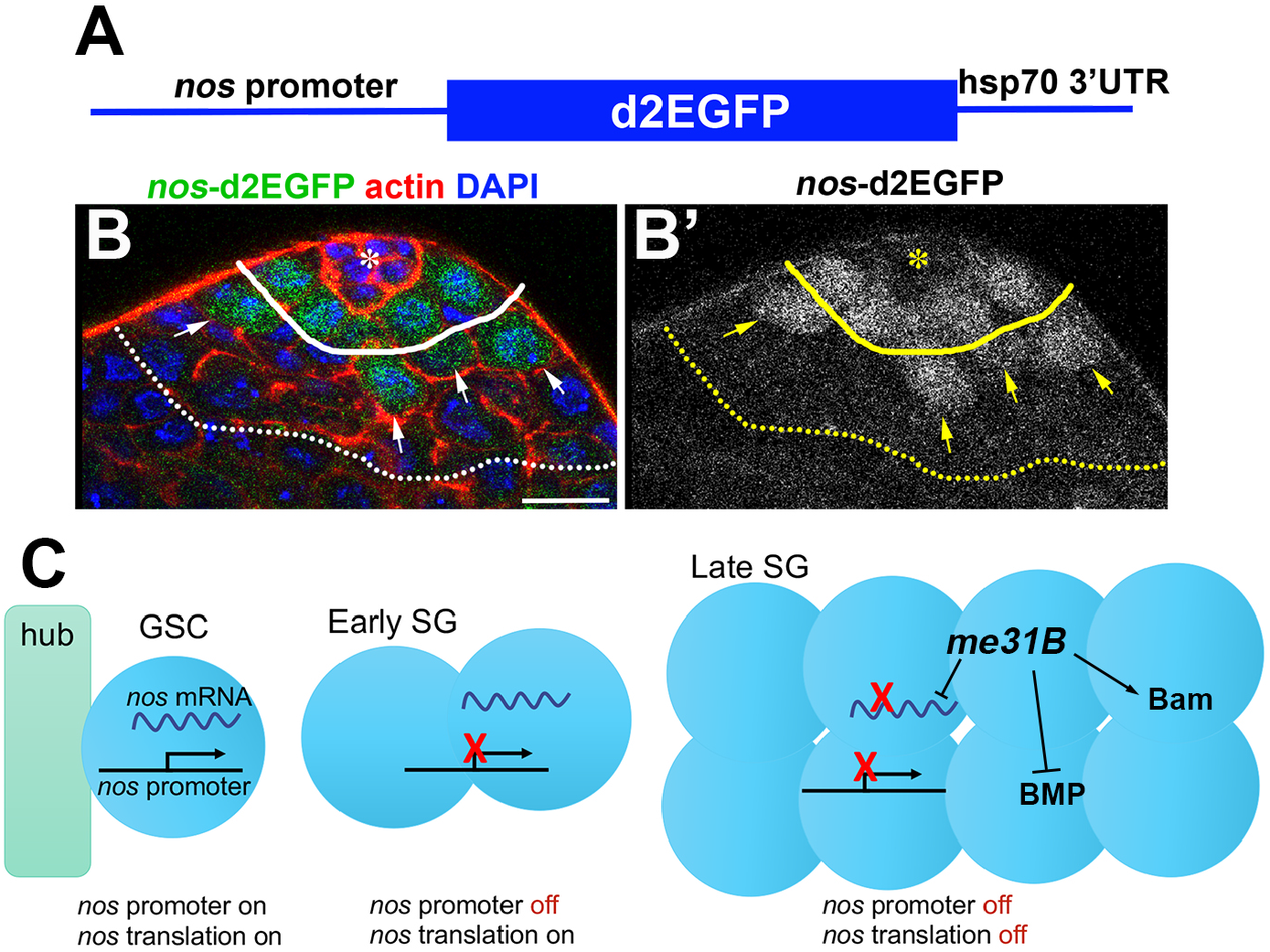
*nos* is transcriptionally and translationally regulated during *Drosophila* spermatogenesis. A. Diagram of *nos* transcription reporter, where *nos* promoter drives unstable GFP protein and 3’UTR sequence from *hsp70*, which makes mRNA short-lived. B. Apical tip of the testis expressing nos transcription reporter. GSC-GB boundary is indicated by solid line, and 4-cell/8-cell SG boundary is indicated by dotted line. GBs that are still connected to GSCs, thus still expressing *nos* transcription reporter are indicated by arrows. Bar: 10μm. Hub is indicated by the asterisk. C. Model of *nos* regulation during germ cell development. In GSCs, the *nos* gene is transcribed and its mRNA is translated, leading to high Nos protein level and thus GSC maintenance. In early SGs, the *nos* gene is no longer transcribed, but Nos protein is produced via translation of inherited *nos* mRNA. In late SGs, the *nos* gene is no longer transcribed, and translation *nos* mRNA is inhibited by *me31B*. This leads to disappearance of Nos protein in these cells, promoting their differentiation.

These results reveal dynamic regulation of *nos* expression through multiple layers (Figure 6C): 1) GSCs and GBs actively transcribe *nos* mRNA, which is translated to produce Nos protein. 2) 2- and 4-cell SGs no longer transcribe *nos* but inherit *nos* mRNA, and thus produce Nos protein. 3) ≥8-cell SGs to not transcribe *nos* mRNA, and translation of inherited *nos* mRNA is inhibited by Me31B, leading to overall downregulation of Nos protein. Loss of Me31B leads to increased translation of *nos* mRNA and increased levels of Nos protein that perdure throughout differentiation, promoting dedifferentiation at later stages.

## Discussion

Stem cell maintenance is critically important for long-term tissue homeostasis. Despite their ability to self-renew, stem cells are not immortal and their life span is often shorter than that of the organism. Dedifferentiation can replenish stem cell pools via conversion of more differentiated cells back into stem cell identity. However, uncontrolled dedifferentiation can lead to tumorigenesis (Landsberg et al., 2012; Schwitalla et al., 2013), thus proper control of dedifferentiation must be essential. Despite its importance, the mechanisms that regulate dedifferentiation are poorly understood.

This study identified *me31B* as a previously unknown and key negative regulator of dedifferentiation through its ability to regulate *nos* mRNAs. Both *nos* and *bam* mRNAs co-immunoprecipitated with Me31B-GFP (Figure 3G). Me31B may reinforce the known antagonistic relationship between *nos* and *bam* germline (Chen and McKearin, 2005; Li et al., 2009) by independently regulating these transcripts (Figure 6C). In addition to extending Nos protein expression to 8-cell SGs and delaying Bam protein expression during germline development, depletion of *me31B* resulted in upregulation of BMP signaling, leading to an increased frequency of dedifferentiating SG cysts (Figure 2). It remains unknown whether *me31B* directly regulates any components of BMP signaling. However, given the antagonistic relationship between *nos* and *bam*, and that BMP signaling represses *bam* expression (Chen and McKearin, 2003a, 2005; Chen and McKearin, 2003b; Harris et al., 2011; Li et al., 2012; Li et al., 2009; Song et al., 2004; Wang and Lin, 2004), it is possible that BMP upregulation can be explained as a downstream effect of misregulated *nos* and/or *bam.*

It remains elusive what controls *me31B* to promote differentiation and/or prevent dedifferentiation. Is *me31B* downregulated by conditions that trigger dedifferentiation? We did not observe any changes in Me31B-GFP protein level or localization when dedifferentiation was artificially induced by transient expression of Bam (not shown). In future studies, it will be of interest to investigate whether Me31B senses niche vacancy (missing GSCs) to trigger dedifferentiation of SGs.

The right balance of differentiation and dedifferentiation must be achieved to ensure maintenance of the stem cell pool, while minimizing the risk of tumorigenesis. The results presented in this study suggest that SGs are in a state of transitioning from stem cell identity to full commitment to differentiation. Whereas GSCs produce Nos protein via *nos* mRNA transcription and its translation, 2- and 4-cell SGs produce Nos protein only via translation of inherited *nos* mRNA. We propose that 2- and 4-cell SGs represent a critical cell population that is not yet fully committed to differentiation, as they still have Nos protein like GSCs, but unlike GSCs they no longer transcribe *nos* (Figure 6C). These SGs may hit a perfect balance of Nos protein that maintains their potential to dedifferentiate into GSCs as necessary, but prevents tumorigenesis by shutting down *nos* transcription. Indeed, 2- and 4-cell SGs are known to be most potent for dedifferentiation (Sheng and Matunis, 2011): although this was speculated to be mostly due to their physical proximity to the hub cells, it is also possible that their ‘Nos production state’ (actively producing Nos protein from inherited mRNA) is more suited for dedifferentiation than later SGs. We propose that stepwise transitions from the stem cell state to the differentiated state are key for maintaining the stem cell pool while preventing tumorigenesis. In summary, the present study provides a new insight into how gradual commitment to differentiation is ensured by transcriptional and translational control of a master regulator.

## Acknowledgements

We thank Bloomington *Drosophila* Stock Center, Developmental Studies Hybridoma Bank, and Drs. Dennis McKearin and Liz Gavis for reagents. We thank the Yamashita lab members and Dr. Angela Anderson (Life Science Editors) for discussion and/or comments on this manuscript. This research was supported by Howard Hughes Medical Institute (to Y.Y) and in part by the NIH Career Training in Reproductive Biology (5T32HD079342-04) (to L.S).

